# A novel reporter allele for monitoring *Dll4* expression within the embryonic and adult mouse

**DOI:** 10.1101/136713

**Authors:** Alexander M. Herman, Alexander M. Rhyner, W. Patrick Devine, Sean P. Marrelli, Benoit G. Bruneau, Joshua D. Wythe

**Author notes:** To whom correspondence should be addressed: Joshua D. Wythe CVRI, Department of Molecular Physiology and Biophysics, Baylor College of Medicine, One Baylor Plaza, Houston, TX 77030.

## Abstract

Canonical Notch signaling requires the presence of a membrane bound ligand and a corresponding transmembrane Notch receptor. Receptor engagement induces multiple proteolytic cleavage events culminating in the nuclear accumulation of the Notch intracellular domain and its binding to a transcriptional co-factor to mediate gene expression. Notch signaling networks are essential regulators of vascular patterning and angiogenesis, as well as myriad other biological processes. *Delta-like 4* (*Dll4*) encodes the earliest Notch ligand detected in arterial cells, and is enriched in sprouting endothelial tip cells. *Dll4* expression has often been inferred by proxy using a *lacZ* knockin reporter allele. This is problematic, as a single copy of *Dll4* is haploinsufficient. Additionally, Notch activity regulates *Dll4* transcription, making it unclear whether these reporter lines accurately reflect *Dll4* expression. Accordingly, accurately defining *Dll4* expression is essential for determining its role in development and disease. To address these limitations, we generated a novel BAC transgenic allele with a nuclear-localized β-galactosidase reporter (*Dll4-BAC-nlacZ*). Through a comparative analysis, we show the BAC line overcomes previous issues of haploinsufficiency, it recapitulates *Dll4* expression *in vivo*, and allows superior visualization and imaging. As such, this novel *Dll4* reporter is an important addition to the growing Notch toolkit.

**Summary Statement:**

We have developed a novel reporter line, free from complicating factors associated with previous alleles, for monitoring *Dll4* expression, at a cellular resolution, in the developing and adult mouse.

## Introduction

Arterial and venous blood vessels are anatomically, functionally, and molecularly distinct. In vertebrates, the proper function of an intact, closed circulatory system requires establishing and maintaining these separate endothelial cell fates. In the vascular system, the Notch signaling pathway is required for proper establishment of arterial and venous endothelial identity (Fischer et al., 2004; Krebs et al., 2004; Krebs et al., 2000; Lawson et al., 2001; Shirayoshi et al., 1997; Shutter et al., 2000; Swiatek et al., 1994; Uyttendaele et al., 2000; Uyttendaele et al., 2001; Uyttendaele et al., 1996; Xue et al., 1999). Specification of endothelium into arteries and veins involves a cascade of signaling events that begin during embryogenesis (Coultas et al., 2005; Fish and Wythe, 2015; Gale and Yancopoulos, 1999).

Current models propose that Sonic Hedgehog activation of the receptor Smoothened induces *Vascular endothelial growth factor* (*Vegf*) transcription (Coultas et al., 2010; Lawson et al., 2002; Vokes et al., 2004). In turn, VEGF activation of VEGF-Receptor 2 (VEGFR2), which is required for arteriovenous specification in the early embryo (Shalaby et al., 1995), initiates expression of *Delta-like 4* (*Dll4*) selectively within arterial endothelial cells. *Dll4* encodes a transmembrane ligand for the Notch family of receptors (Shutter et al., 2000). Notch1, as well as its essential transcriptional co-factor Rbpj-k (also known as CSL, Su(H), CBF) are essential regulators of arteriovenous patterning in the early vertebrate embryo, as their deletion leads to arteriovenous malformations and embryonic lethality (Krebs et al., 2004; Krebs et al., 2000).

*Dll4* is a critical regulator of vascular morphogenesis, as its loss results in vascular defects and embryonic lethality by E10.5 (Duarte et al., 2004; Gale et al., 2004; Krebs et al., 2004). *In situ* hybridization results show that *Dll4* is the earliest Notch ligand detected in arterial precursor cells (aPCs), potentially preceding expression of Notch receptors (Chong et al., 2011; Lindskog et al., 2014; Mailhos et al., 2001; Wythe et al., 2013). Unlike *Notch1, Dll4* expression in the dorsal aorta does not require hemodynamic force in the early mouse embryo, and is invariably arterial specific (Chong et al., 2011; Jahnsen et al., 2015). Conversely, Dll4 and Notch gain-of-function manipulations alter arteriovenous patterning and lead to lethality with obvious AV patterning defects in embryos (Kim et al., 2008; Krebs et al., 2004; Trindade et al., 2008; Wythe et al., 2013), and AVMs in adults (Carlson et al., 2005; Murphy et al., 2014; Murphy et al., 2008).

In addition to regulating AV specification, Dll4 function also controls angiogenesis. The dynamic expression of *Dll4* within the tip cell, and its repression in the trailing stalk cells that make up a sprouting vessel is controlled by VEGF-VEGFR2 signaling (Gerhardt et al., 2003; Hellstrom et al., 2007; Lobov et al., 2007). Dll4-Notch signaling acts as a negative feedback regulator of VEGFR2 to establish the proper ratio of tip to stalk cells in the sprouting vasculature. Consequently, loss of *Dll4*, or *Rbpj-k*, leads to increased endothelial proliferation and hypersprouting (Jakobsson et al., 2010; Suchting et al., 2007).

Molecular and biochemical methods to query *Dll4* expression, such as *in situ* hybridization, or antibody-based immunostaining, can be time consuming, and yield variable results. Mouse models with a *lacZ* reporter cassette replacing the translational start site of endogenous *Dll4* have been used to visualize *Dll4* expression, however these modifications create a null allele (Duarte et al., 2004; Gale et al., 2004). In the case of *Dll4* this is problematic, as these two lines, as well as a third, conventional loss of function allele (Krebs et al., 2004), demonstrated that heterozygous *Dll4* mutants displayed incompletely penetrant, lethal haploinsufficiency between E9.5 and E10.5 (Duarte et al., 2004; Gale et al., 2004; Krebs et al., 2004). Outcrossing these lines to different genetic backgrounds reduces the penetrance of this effect, but the ratio of viable offspring remains low (Benedito and Duarte, 2005; Duarte et al., 2004). Furthermore, interpreting *Dll4* expression levels in these knockin/knockout reporter mice is complicated due to a positive feedback loop between *Dll4* expression and Notch signaling (Caolo et al., 2010). As such, even in viable mutant animals, it is not clear if the knockin reporter faithfully recapitulates *Dll4* expression. Precisely defining *Dll4* expression in the embryo and adult is central to understanding its role during vascular specification, angiogenesis (Hellstrom et al., 2007), T-cell development (Koch et al., 2008), and retinogenesis (Luo et al., 2012). Finally, Dll4 may signal to Notch receptors in even more tissues, such as the gut or kidney (Benedito and Duarte, 2005), necessitating an accurate, reliable, and robust method for visualizing its expression domain *in vivo*.

In the case of *Dll4*, histochemical detection of β-gal is considered more sensitive than detection of *Dll4* mRNA by *in situ* hybridization (Benedito and Duarte, 2005). To retain this advantage, but overcome the inherit drawbacks of available *Dll4* reporter-knockout mouse lines, we generated a transgenic *Dll4-BAC-nlacZ* reporter line. Herein we show that this line faithfully recapitulates endogenous *Dll4* expression in the embryonic, postnatal, and adult mouse, while avoiding potential confounds associated with disrupted Notch signaling. Furthermore, the signal strength in this model is greater than previous *Dll4* reporter lines, and addition of a nuclear localization signal increases cellular resolution. Going forward, this novel tool will facilitate studies of *Dll4* expression within the embryonic and adult mouse.

## Results

Using recombineering, a nuclear localized *lacZ* reporter cDNA cassette (*nlacZ*) was targeted to the start codon of murine *Dll4* in a bacterial artificial chromosome (BAC) to generate a *Dll4* reporter construct (Fig. 1A) (Warming et al., 2005). The BAC clone, spanning approximately 81 kb of mouse chromosome 2, contained the entire *Dll4* locus, as well as approximately 32.5 kb upstream and 38 kb downstream. The full-length, recombined clone, *Dll4-BAC-nlacZ,* was linearized and used to create transgenic mice by pronuclear injection. From one round of injections, 2 successful founders (*Dll4-BAC-nlacZ*^4336^ and *Dll4-BAC-nlacZ*^4316^) were identified with germ line transmission of the transgene. We focused our studies on founder *Dll4-BAC-nlacZ*^4336^.

**Figure 1:**
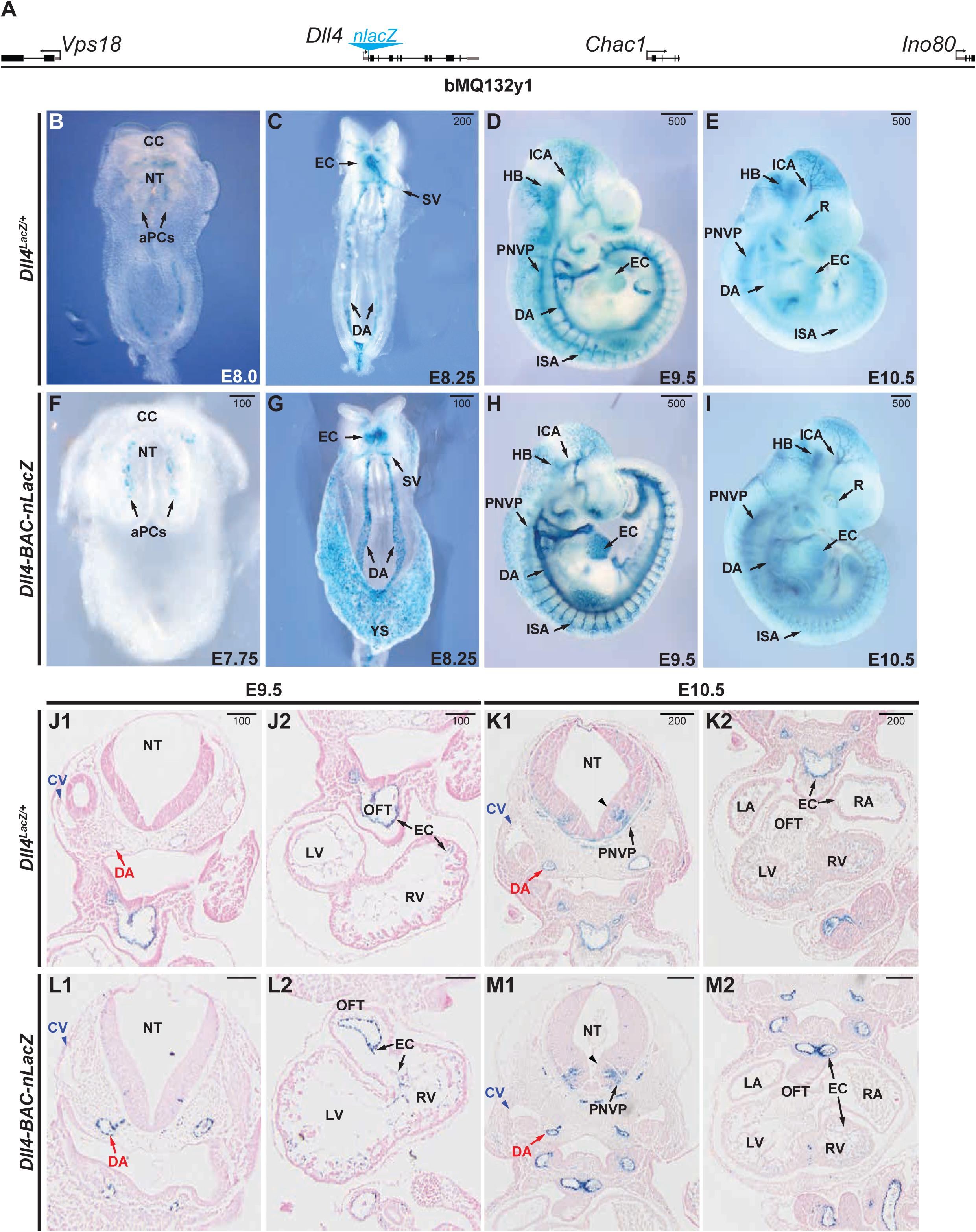
Comparative *Dll4* expression during early embryonic development. **A,** Schematic of the BAC transgene used for generating the *Dll4-BAC-nlacZ* mouse line. **B-E,** β-gal activity in E7.75 - E10.5 *Dll4*^*lacZ/+*^ mouse embryos, ventral (B, C) and sagittal (D, E) views. **F-I,** β-gal activity in E7.75 - E10.5 *Dll4-BAC-nlacZ* mouse embryos, ventral (F, G) and sagittal (H, I) views. **J-M,** Coronal view of X-gal-stained and eosin-counterstained sections of E9.5 and E10.5 *Dll4*^*lacZ/+*^ embryos. **N-Q,** Coronal view of X-gal-stained and eosin-counterstained sections of E9.5 and E10.5 *Dll4-BAC-nlacZ* embryos. Abbreviations: aPCs – aortic progenitor cells; CC – cardiac crescent; CV – cardinal vein; DA – dorsal aorta; EC – endocardium, ICA – internal carotid artery, ISA – intersegmental artery; LA – left atrium; LV – left ventricle; NT – neural tube; OFT – outflow tract; PNVP – perineural vascular plexus; RA – right atrium; RV – right ventricle; SV – sinus venosus; caret – ventral V2 interneuron population.

To validate our BAC transgenic line, we compared its β-galactosidase (β-gal) activity to that of *Dll4*^*lacZ/+*^ in the embryonic and postnatal mouse at different developmental time points (Fig. 1B-I). Prior work has suggested *Dll4* transcripts are initiated at E8.0 (Benedito and Duarte, 2005), however, using the same allele employed in that study, as well as our novel BAC line, we detect β-gal at E7.75 in the presumptive endocardium of the cardiac crescent, as well as in aortic progenitor cells (aPCs) (Fig. 1B and 1F), in agreement with previous reports examining endogenous *Dll4* transcripts (Benedito and Duarte, 2005; Duarte et al., 2004; Mailhos et al., 2001; Shutter et al., 2000; Wythe et al., 2013). By E8.25, analogous to *Dll4* mRNA (Chong et al., 2011; Wythe et al., 2013), *lacZ* expression was present in the endocardium and sinus venosus, as well as the dorsal aorta (Fig. 1C and 1G). By E9.5, the dorsal aorta, endocardium, internal carotid artery, hindbrain, intersomitic arterial vessels, and perineural vascular plexus all displayed β-gal activity (Fig. 1D and 1H). At E10.5, both reporters labelled each of these structures, as well as the retina (Fig. 1E and 1I). Histological analyses of E9.5 and E10.5 embryos revealed that while β-gal was observed in the dorsal aorta and endocardium of both *Dll4*^*lacZ/+*^ and *Dll4-BAC-nlacZ* mice, it was absent from the cardinal vein (Fig. 1J1-M2), confirming its arterial specificity within the endothelium. Notably, at E10.5, *lacZ* was expressed within a narrow, ventral stripe of tissue in the neural tube (presumably V2 interneurons), in agreement with previous reports (Benedito and Duarte, 2005; Mailhos et al., 2001).

*Dll4*^*lacZ/+*^ mutants can exhibit developmental delayed (Duarte et al., 2004). *Dll4-BAC-nlacZ* mice were normal in size at all stages examined. This difference became more apparent during later embryogenesis (E12.5, E14.5, and E18.5) (Fig. 2A-F), although the domain of *lacZ* expression at the wholemount level was comparable between the two lines. By E14.5, superficial β-gal activity within the skin was evident in both lines (Fig. 2B and 2E), but it was not clear if *Dll4* reporter activity was restricted to the arterial endothelium, or present within other vessel types, such as the venous vasculature or lymphatic system, as suggested by previous reports (Bernier-Latmani et al., 2015; Niessen et al., 2011). To determine the identity of β-gal-positive cells, skin from the E14.5 forelimb of both genotypes was processed for immunohistochemistry using antibodies against β-gal, the endothelial-specific cell surface receptor CD31 (PECAM), the lymphatic vessel-specific antigen Podoplanin, the arterial-specific smooth muscle cell protein alpha smooth muscle actin (SMA), the neuronal-specific marker Tuj1, as well as endogenous Dll4. Confocal microscopy revealed that β-gal-positive cells of either genotype did not colocalize with Podoplanin or Tuj1, but did colocalize with CD31, SMA, and Dll4 (Fig. 3A1-H6), demonstrating that *lacZ* expression was restricted to the arterial vasculature.

**Figure 2:**
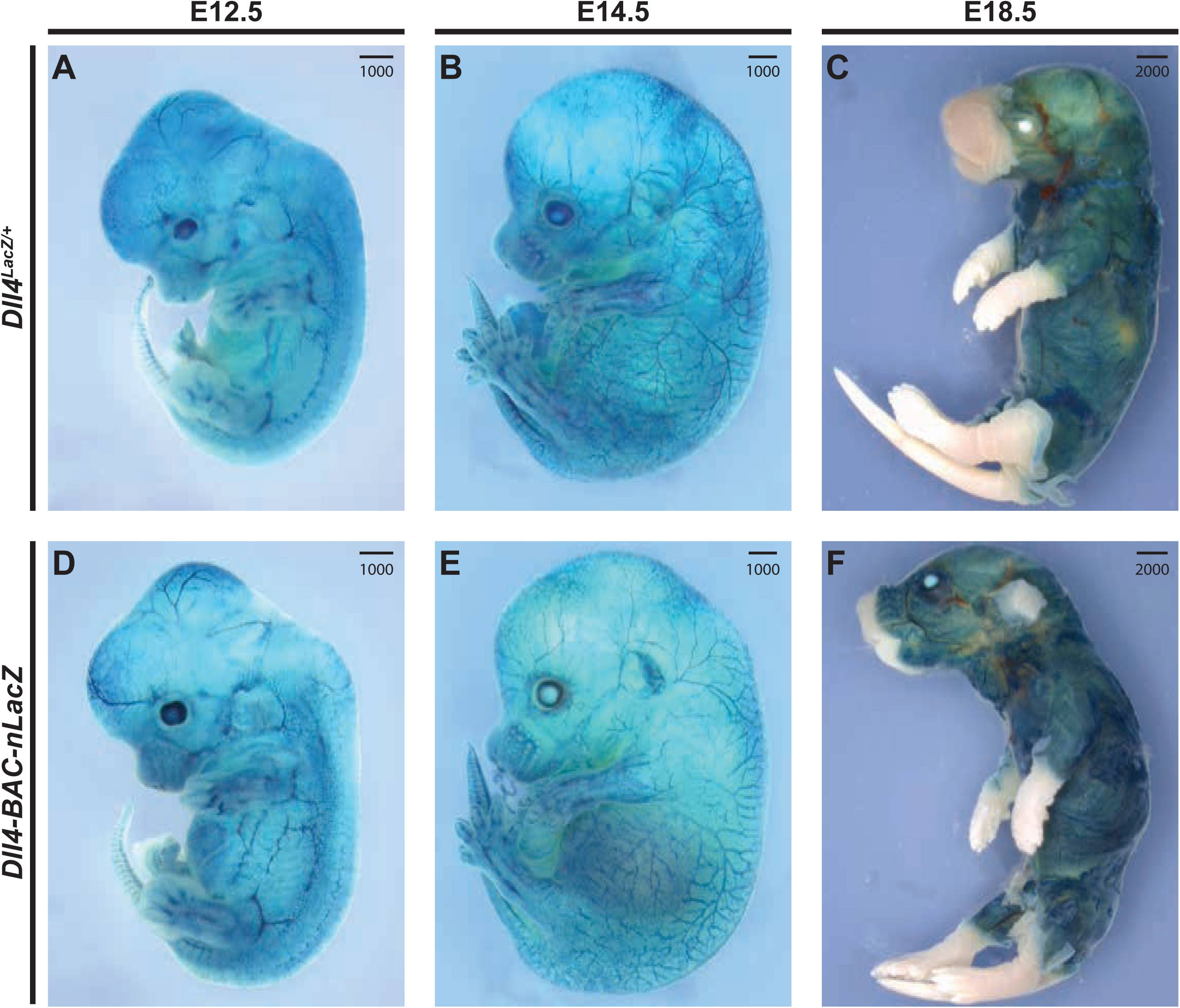
Comparative *Dll4* expression during intermediate and late-stage embryonic development. **A-C,** lacZ activity in E12.5 (A), E14.5 (B), and E18.5 (C) *Dll4*^*lacZ/+*^ mouse embryos. **D-F,** lacZ activity in E12.5 (D), E14.5 (E), and E18.5 (F) *Dll4-BAC-nlacZ* mouse embryos. Noticeable size differences can be observed between genotypes due to heterozygous *Dll4* loss of function.

**Figure 3:**
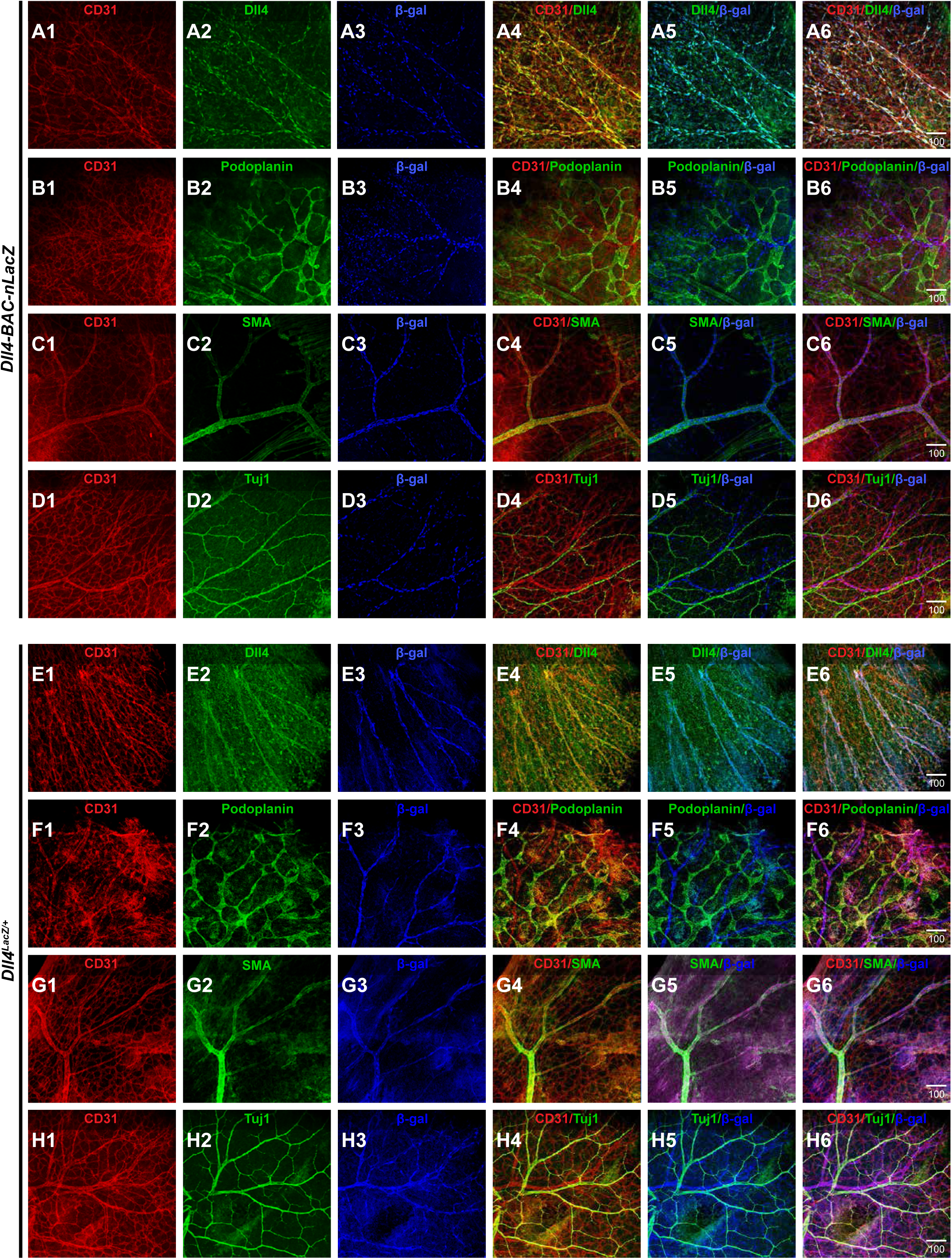
β-gal reporter activity is restricted to arterial vasculature in the skin. **A1-A6,** Single channel views of indirect immunofluorescence for CD31 (A1), Dll4 (A2), β-gal (A3), and merged (A4-A6) images showing colocalization between β-gal-positive vasculature and endogenous Dll4 in *Dll4-BAC-nlacZ* mouse skin. **B1-B6,** CD31 (B1), Podoplanin (B2), β-gal (B3), and merged (B4-B6) images showing a lack of colocalization between β-gal-positive vasculature and the lymphatic-specific marker Podoplanin in *Dll4-BAC-nlacZ* mouse skin. **C1-C6,** CD31 (C1), SMA (C2), β-gal (C3), and merged (C4-C6) images showing colocalization between β-gal-positive vasculature and the arterial-specific marker, smooth muscle actin (SMA). **D1-D6,** CD31 (D1), Tuj1 (D2), β-gal (D3), and merged (D4-D6) images showing lack of colocalization between β-gal-positive vasculature and the neuronal-specific marker Tuj1. **E1-E6,** CD31 (E1), Dll4 (E2), β-gal (E3), and merged (E4-E6) images showing colocalization between β-gal-positive vasculature and endogenous Dll4 in *Dll4*^*lacZ/+*^ mouse skin. **F1-F6,** CD31 (F1), Podoplanin (F2), β-gal (F3), and merged (F4-F6) images showing a lack of colocalization between β-gal-positive vasculature and the lymphatic-specific marker Podoplanin in *Dll4*^*lacZ/+*^ mouse skin. **G1-G6,** CD31 (G1), SMA (G2), β-gal (G3), and merged (G4-G6) images showing colocalization between β-gal-positive vasculature and the arterial-specific marker SMA. **H1-H6,** CD31 (H1), Tuj1 (H2), β-gal (H3), and merged (H4-H6) images showing lack of colocalization between β-gal-positive vasculature and the neuronal-specific marker Tuj1.

At these same embryonic stages, the brains (Fig. 4), hearts (Fig. 5), and lungs (Fig. 5) from the endogenous knockin and BAC transgenic reporter embryos were examined and compared. Within the embryonic brain, the expression domains of *Dll4*^*lacZ/+*^ and *Dll4-BAC-nlacZ* were almost indistinguishable from E12.5 through E18.5 at the wholemount level, with signal evident within the vertebral arteries (VA), basilar artery (BA), superior cerebellar arteries (SCA), posterior cerebral arteries (PCA), middle cerebral arteries (MCA), and anterior cerebral arteries (ACA), as well as their respective branches. Collaterals linking the MCA, ACA, and PCA territories became evident between E14.5 and E18.5, consistent with previous reports (Chalothorn and Faber, 2010). Histological analysis revealed reporter activity throughout the brains in both lines, from the olfactory bulb to the brain stem, and the cortex to the hypothalamus (Fig. 4).

**Figure 4:**
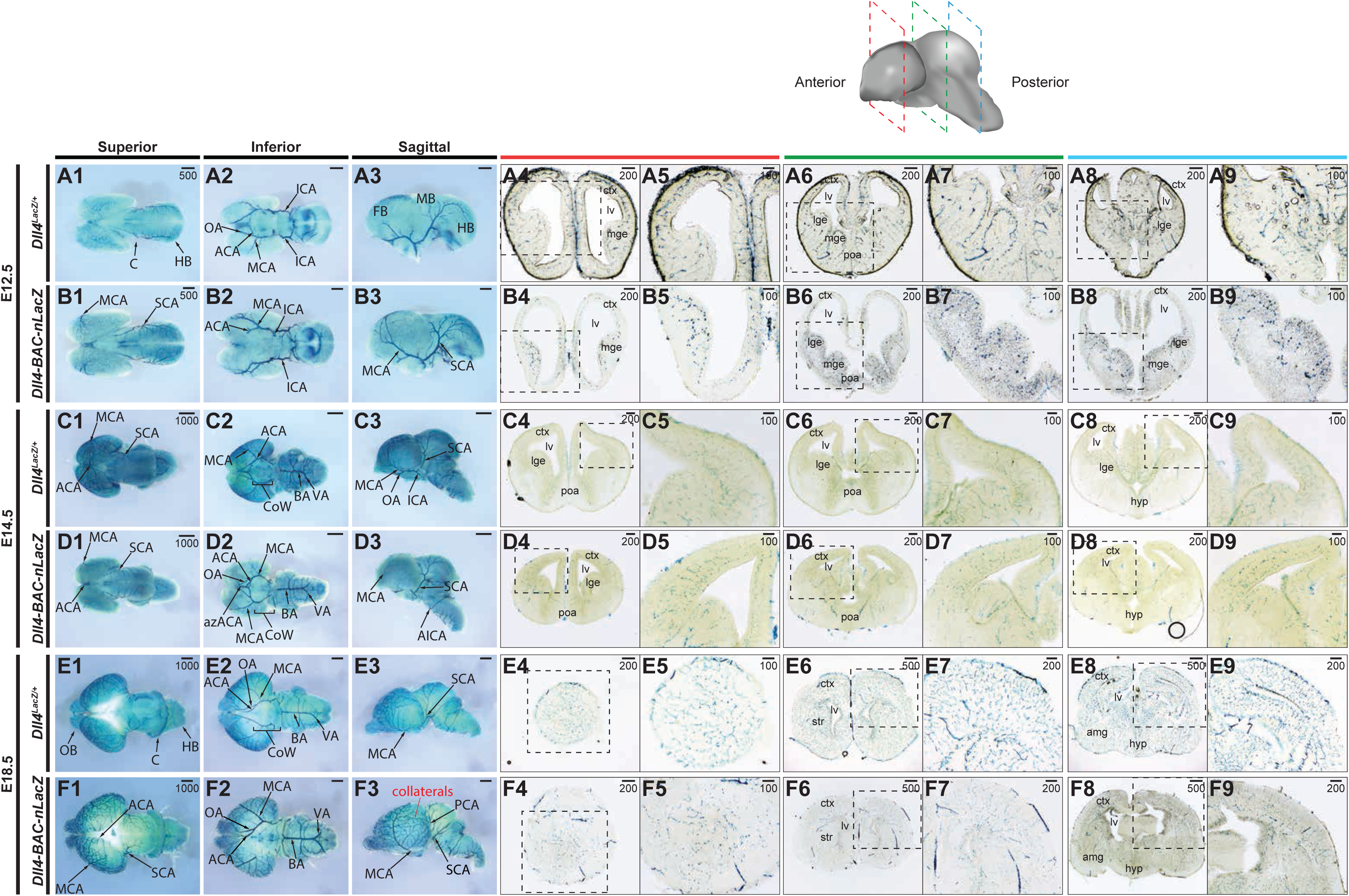
Comparative *Dll4* expression in intermediate and late-stage wholemount embryonic brains. **A-B,** β-gal activity in the E12.5 embryonic brain of (A) *Dll4*^*lacZ/+*^ and (B) *Dll4-BAC-nlacZ* mice. **A1-A3** and **B1-B3** show representative wholemount images of the brain from superior, inferior, and sagittal planes. **A4-A9** and **B4-B9** show representative coronal sections through the brain, from anterior to posterior. **C-D,** β-gal activity in the E14.5 embryonic brain of (C) *Dll4*^*lacZ/+*^ and (D) *Dll4-BAC-nlacZ* mice. **C1-C3** and **D1-D3** show representative wholemount images of the brain from superior, inferior, and sagittal planes. **C4-C9** and **D4-D9** show representative coronal sections through the brain, from anterior to posterior. **E-F,** β-gal activity in the E18.5 embryonic brain of (E) *Dll4*^*lacZ/+*^ and (F) *Dll4-BAC-nlacZ* mice. **E1-E3** and **F1-F3** show representative wholemount images of the brain from superior, inferior, and sagittal planes. **E4-E9** and **F4-F9** show representative coronal sections through the brain, from anterior to posterior. Abbreviations: ACA – anterior cerebral artery; azACA – azygos of the anterior cerebral artery; MCA – middle cerebral artery; SCA – superior cerebellar artery; AIC – anterior inferior cerebellar artery; VA – vertebral artery; ICA – internal carotid artery; BA – basilar artery; PCA – posterior cerebral artery.

**Figure 5:**
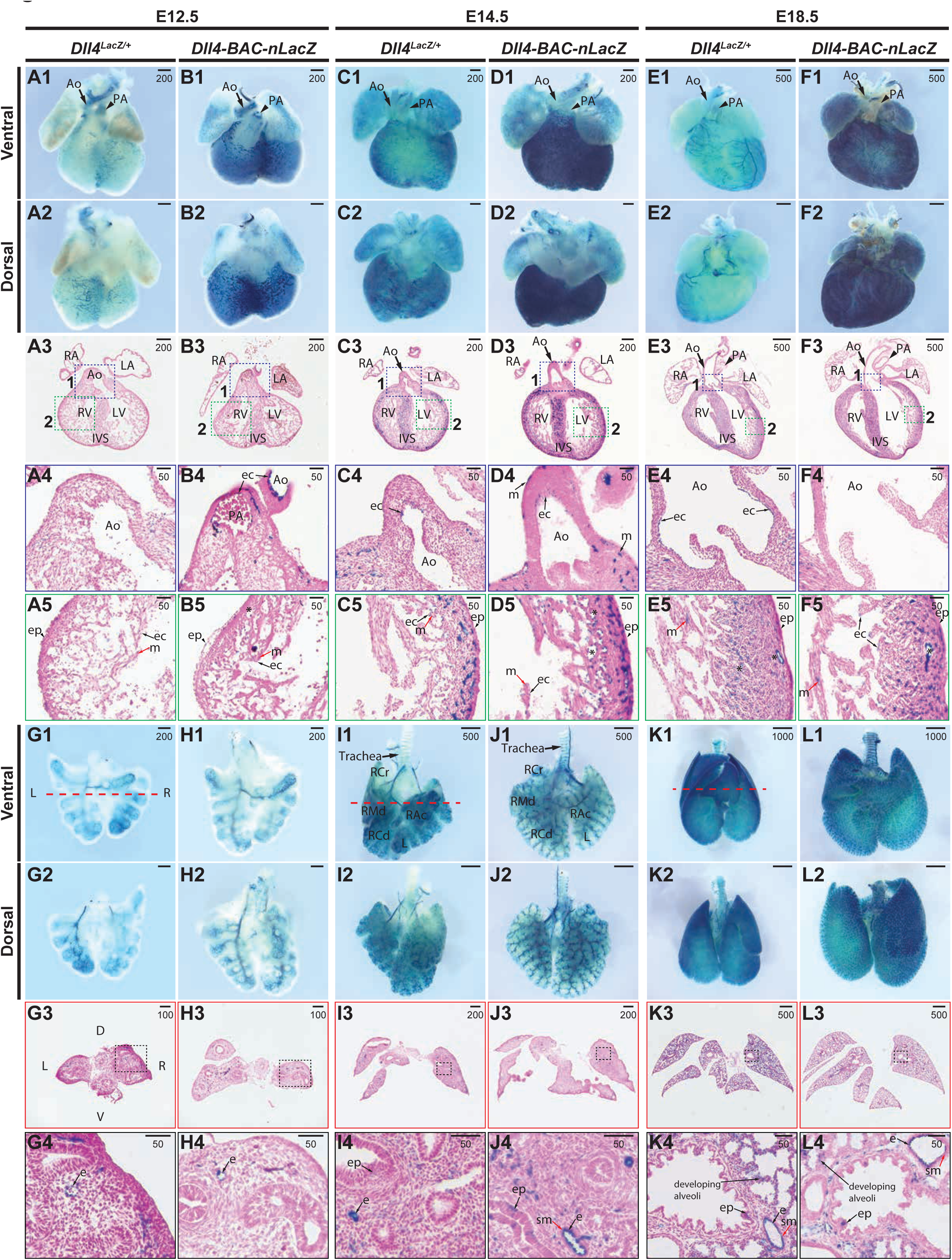
Comparative *Dll4* expression in intermediate and late-stage embryonic hearts and lungs. **A-B,** β-gal activity in E12.5 hearts from (A) *Dll4*^*lacZ/+*^ or (B) *Dll4-BAC-nlacZ* mice. **A1-A2** and **B1-B2** show representative wholemount hearts from *Dll4*^*lacZ/+*^ and *Dll4-BAC-nlacZ* mice, respectively, from ventral and dorsal views. **A3** and **B3** show β-gal activity in a representative cross-section through the heart, which is magnified accordingly in panels **A4-A5** and **B4-B5.** β-gal activity is present within coronary plexus (A1, A2, B1, B2), the endocardium of the distal end (A1, B1) and root of the aorta (A4, B4) and pulmonary artery in both lines, as well as the endocardium and subepicardial vasculature (A5, B5), but absent from the epicardium and myocardium. **C-D,** β-gal activity in E14.5 hearts from (C) *Dll4*^*lacZ/+*^ or (D) *Dll4-BAC-nlacZ* mice. **C1-C2** and **D1-D2** show representative wholemount hearts from *Dll4*^*lacZ/+*^ and *Dll4-BAC-nlacZ* mice, respectively, from ventral and dorsal views. **C3** and **D3** show β-gal activity in a representative cross-section through the heart, which is magnified accordingly in panels **C4-C5** and **D4-D5.** β-gal activity is localized to the endocardium of the aorta in both lines (C4, D4), as well as the chamber endocardium (C5, D5), and subepicardial coronary vasculature (C5, D5). β-gal activity was also detected within a small fraction of the myocardium in the BAC reporter line at this stage. **E-F,** β-gal activity in E18.5 hearts from (E) *Dll4*^*lacZ/+*^ or (F) *Dll4-BAC-nlacZ* mice. **E1-E2** and **F1-F2** show representative wholemount hearts from *Dll4*^*lacZ/+*^ and *Dll4-BAC-nlacZ* mice, respectively, from ventral and dorsal views. **E3** and **F3** show β-gal activity in a representative cross-section through the heart, which is magnified accordingly in panels **E4-E5** and **F4-F5.** β-gal activity is localized to the endocardium of the aortic root *Dll4*^*lacZ/+*^ but absent from *Dll4-BAC-nlacZ* mice (E4, F4), and present in both lines within the chamber endocardium and coronary vasculature (E5, F5), and sparsely in the myocardium. Abbreviations: Ao – aorta; ec – endocardium; ep – epicardium; m – myocardium; IVS – interventricular septum; LA – left atrium; LV – left ventricle; PA – pulmonary artery; RA – right atrium; RV – right ventricle. **G-H,** β-gal activity in E12.5 lungs from (A) *Dll4*^*lacZ/+*^ or (B) *Dll4-BAC-nlacZ* mice. **G1-G2** and **H1-H2** show representative wholemount lungs from *Dll4*^*lacZ/+*^ and *Dll4-BAC-nlacZ* mice, respectively, from ventral and dorsal views. **G3** and **H3** are representative cross-sections through the lungs, and boxed in areas are magnified in **G4** and **H4**, revealing activity within the endothelium**. I-J,** β-gal activity in E14.5 lungs from (C) *Dll4*^*lacZ/+*^ or (D) *Dll4-BAC-nlacZ* mice. **I1-I2** and **J1-J2** show representative wholemount lungs from *Dll4*^*lacZ/+*^ and *Dll4-BAC-nlacZ* mice, respectively, from ventral and dorsal views. **I3** and **J3** show β-gal activity in a representative cross-section through the lungs, which is magnified accordingly in panels **I4** and **J4**, shows endothelial-specific activity in both lines. **K-L,** β-gal activity in E18.5 lungs from (E) *Dll4*^*lacZ/+*^ or (F) *Dll4-BAC-nlacZ* mice. **K1-K2** and **L1-L2** show representative wholemount lungs from *Dll4*^*lacZ/+*^ and *Dll4-BAC-nlacZ* mice, respectively, from ventral and dorsal views. **K3** and **L3** show β-gal activity in a representative cross-section through the lungs, which is magnified accordingly in panels **K4** and **L4**, shows endothelial-specific activity in both lines. Abbreviations: L – left; R – right; V – ventral; D – dorsal; e – endothelium; sm – smooth muscle.

At E12.5, *Dll4*^*lacZ/+*^ activity was evident at the wholemount level within the great vessels (aorta and pulmonary artery) and the primary plexus that ultimately generate the coronary vasculature of the embryonic heart (Fig. 5 A1-A2). Histological analysis confirmed that β-gal was restricted to the endothelial lining of the great vessels, as well as the endocardium of the atrial and ventricular chambers, the endothelium underlying the epicardium, and vessels within the compact myocardium (Fig. 5A3-A5). This same pattern of activity was observed in *Dll4-BAC-nlacZ* hearts (Fig. 5B1-B5), although labeling of the primary coronary plexus was more robust in the BAC reporter line. At E14.5, in *Dll4*^*lacZ/+*^ embryos, reporter signal was evident within the great vessels and the atria, as well as the coronary plexus (Fig. 5C1-C5). *Dll4-BAC-nlacZ* activity was present in a similar domain, with signal present throughout the endothelium lining the great vessels, the chamber endocardium, and the coronary vessels underlying the compact myocardium (Fig. 5D1-D5). At E18.5, *Dll4*^*lacZ/+*^ expression persisted within the aorta, pulmonary artery, and coronary vessels (Fig. 5E1-E5). Activity was also detected within the freewall myocardium (Fig. 5E5). At this stage *Dll4-BAC-nlacZ* signal was diminished within the endothelium of the pulmonary artery, and was not detected within the aortic root (Fig. 5F1-F4). β-gal was also detected in both the endocardium and myocardium of the atrial and ventricular chambers (Fig. 5F3 and F5).

We next examined reporter activity within the embryonic lung (Fig. 5G-L). Similar to the heart (Fig. 5A-F), lungs were noticeably smaller in *Dll4*^*lacZ/+*^ animals compared to the BAC reporters. In the pseudoglandular stage (E12.5), β-gal activity was detected within the primitive vascular tree of the left and right lobe in *Dll4*^*lacZ/+*^ animals, with no discernable difference in expression from the rostral to caudal axis, or in any of the lobes (Fig. 5G1-G2). Histological analysis revealed signal within the endothelium of large and small caliber vessels (Fig. 5G1-G4). This expression pattern was recapitulated in *Dll4-BAC-nlacZ* animals (Fig. 5H1-H4). By the canalicular stage (E14.5), β-gal activity was present in narrow, horizontal bands across the ventral side of the trachea in both the knockin and BAC animals, as well as the vascular tree and endothelium, but excluded from smooth muscle (Fig. 5I1-J4). Interestingly, expression was observed —albeit infrequently— within the airway epithelium in both the knockin and the BAC lines (Fig. 5I4, J4). At the saccular stage (E18.5), β-gal activity within the trachea and vascular tree persisted in both samples, but was more evident in *Dll4-BAC-nlacZ* animals (Fig. 5K1-L4). X-gal reactivity was infrequently detected in the airway epithelium of either line, but present within the developing distal alveoli of both reporters, presumably in the capillary endothelium (Fig. 5K4, L4). In both lines, signal was evident in the endothelium of small and medium size vessels, but absent in smooth muscle (Fig. 5K4, L4).

Across all embryonic tissues examined (brain, heart, lung, skin), signal strength and resolution were superior in *Dll4-BAC-nlacZ* animals compared to *Dll4*^*lacZ/+*^ mice, particularly in sectioned tissue, where distinct cells could be observed in *Dll4-BAC-nlacZ* tissue, due to its nuclear localization. Additionally, X-gal staining proceeded more rapidly in *Dll4-BAC-nlacZ* tissue compared to age-matched *Dll4*^*lacZ*^ tissue, in both wholemount and sectioned samples. Overt growth deficits were not observed in embryos derived from either BAC founder line (Figs. 1 and 2, data not shown).

We next surveyed reporter activity in early postnatal and adult tissues. Like the embryonic cranial comparisons, between *Dll4-BAC-nlacZ* and *Dll4*^*lacZ/+*^ cranial regions, within the postnatal and adult brain β-gal labelled the major cerebral arteries, as well as their branches and collaterals in both lines (Fig. 6). Reporter activity spanned the anterior-posterior and dorsal-ventral axes in both lines. Staining of P1 and P5 brains showed increased vessel density and branching of pial arteries in *Dll4-BAC-nlacZ* samples (Fig. 6A and B). Staining in the adult brains appeared grossly similar between the two lines at the wholemount level (Fig. 6E and F). Histological analysis revealed activity throughout the olfactory bulb, cerebral cortex, hippocampus, and cerebellum, in an indistinguishable manner between the two reporters (Fig. 6).

**Figure 6:**
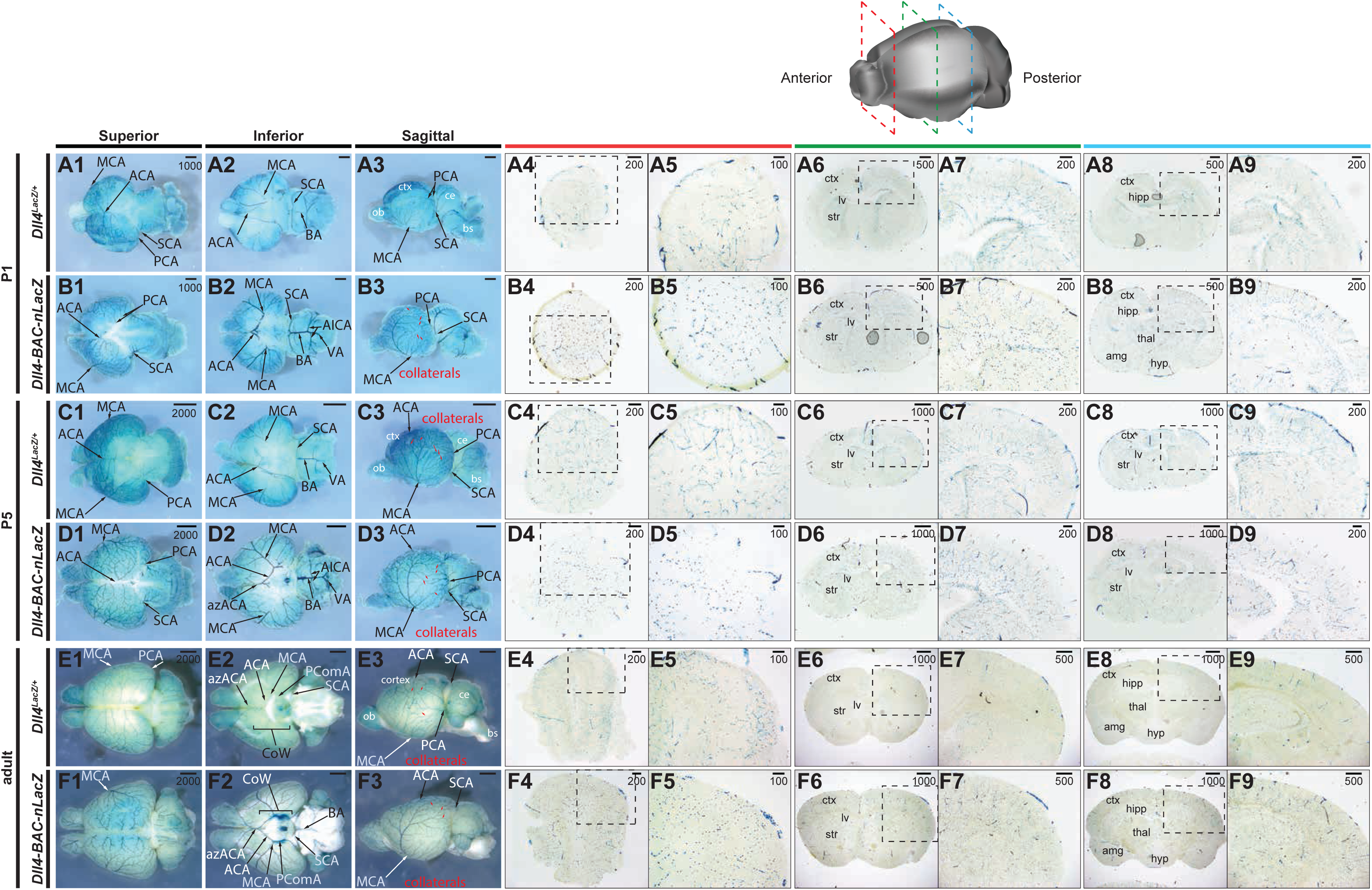
Comparative *Dll4* expression in wholemount postnatal and adult brains. **A-B,** β-gal activity in the P1 postnatal brain of (A) *Dll4*^*lacZ/+*^ and (B) *Dll4-BAC-nlacZ* mice. **A1-A3** and **B1-B3** show representative wholemount images of the brain from superior, inferior, and sagittal planes. **A4-A9** and **B4-B9** show representative coronal sections through the brain, from anterior to posterior. **C-D,** β-gal activity in the P5 postnatal brain of (C) *Dll4*^*lacZ/+*^ and (D) *Dll4-BAC-nlacZ* mice. **C1-C3** and **D1-D3** show representative wholemount images of the brain from superior, inferior, and sagittal planes. **C4-C9** and **D4-D9** show representative coronal sections through the brain, from anterior to posterior. **E-F,** β-gal activity in the adult brain of (E) *Dll4*^*lacZ/+*^ and (F) *Dll4-BAC-nlacZ* mice. **E1-E3** and **F1-F3** show representative wholemount images of the brain from superior, inferior, and sagittal planes. **E4-E9** and **F4-F9** show representative coronal sections through the brain, from anterior to posterior. Abbreviations: ACA – anterior cerebral artery; azACA – azygos of the anterior cerebral artery; MCA – middle cerebral artery; SCA – superior cerebellar artery; AIC – anterior inferior cerebellar artery; VA – vertebral artery; ICA – internal carotid artery; BA – basilar artery; PCA – posterior cerebral artery.

In the P1 postnatal *Dll4*^*lacZ/+*^ wholemount heart, β-gal was active within the coronary vessels, aorta, and pulmonary artery (Fig. 7A1-A2). Sections revealed signal within the chamber endocardium (Fig. 7A3-A5), as well as the endothelial lining of the aorta (Fig. 7A4), the coronary vascular endothelium, and the myocardium (Fig. 7A5). Here, the activity and domain of the *Dll4-BAC-nlacZ* line differed dramatically from the knockin, in that while expression was also detected (sparsely) within the endothelial lining of the aortic root, it robustly labelled the chamber endocardium, myocardium, and the coronary vasculature (Fig. 7B1-B5). At P5, *Dll4*^*lacZ/+*^ drove β-gal within the endothelium of the aorta and pulmonary artery (Fig. 7C1-4), the chamber endocardium, myocardium, and coronary vasculature (Fig. 7C5). The BAC reporter marked these same expression domains, but demonstrated elevated β-gal activity within the myocardium compared to the knockin line (Fig. 7D1-D5). In the adult heart, both lines showed weak signal within the endothelial lining of the aorta, as well as the chamber endocardium, myocardium, and coronary vasculature (Fig. 7E-F). Expression was detected within the epicardium in knockin animals only at P5 (Fig. 7D5), and within BAC reporters only at P1 (Fig. 7B5).

**Figure 7:**
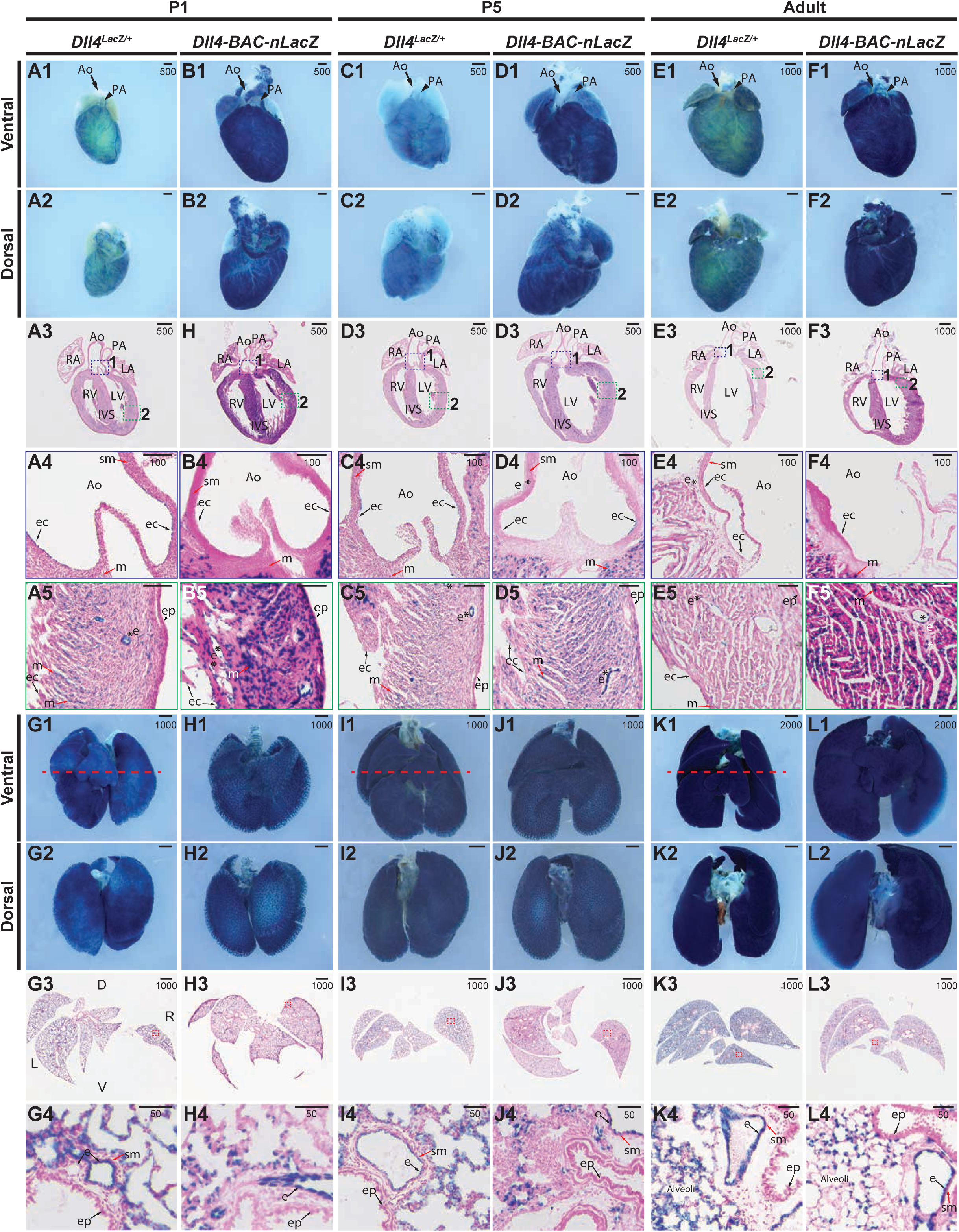
Comparative *Dll4* expression in postnatal and adult hearts and lungs. **A-B,** β-gal activity in P1 hearts from (A) *Dll4*^*lacZ/+*^ or (B) *Dll4-BAC-nlacZ* mice. **A1-A2** and **B1-B2** show representative wholemount hearts from *Dll4*^*lacZ/+*^ and *Dll4-BAC-nlacZ* mice, respectively, from ventral and dorsal views. **A3** and **B3** show β-gal activity in a representative cross-section through the heart, which is magnified accordingly in panels **A4-A5** and **B4-B5**, with activity evident within the endocardial lining of the aorta in both lines (A4, B4), as well as the endocardium, coronary vasculature, and myocardium (A5, B5)**. C-D,** β-gal activity in P5 hearts from (C) *Dll4*^*lacZ/+*^ or (D) *Dll4-BAC-nlacZ* mice. **C1-C2** and **D1-D2** show representative wholemount hearts from *Dll4*^*lacZ/+*^ and *Dll4-BAC-nlacZ* mice, respectively, from ventral and dorsal views. **C3** and **D3** show β-gal activity in a representative cross-section through the heart, which is magnified accordingly in panels **C4-C5** and **D4-D5**, with activity evident within the endocardial lining of the aorta in both lines, with activity persisting in the endocardium of the aorta (C4, D4) and chambers, as well as the myocardium and coronary vasculature (C5, D5). **E-F,** β-gal activity in adult hearts from (E) *Dll4*^*lacZ/+*^ or (F) *Dll4-BAC-nlacZ* mice. **E1-E2** and **F1-F2** show representative wholemount hearts from adult *Dll4*^*lacZ/+*^ and *Dll4-BAC-nlacZ* mice, respectively, from ventral and dorsal views. **E3** and **F3** show β-gal activity in a representative cross-section through the heart, magnified in panels **E4-E5** and **F4-F5.** β-gal activity is localized to the endocardium of the aortic root *Dll4*^*lacZ/+*^ but absent from *Dll4-BAC-nlacZ* mice (E4, F4), and present in both lines within the chamber endocardium and coronary vasculature (E5, F5) (asterisks), and within the myocardium. Abbreviations: Ao – aorta; ec – endocardium; ep – epicardium; m – myocardium; sm – smooth muscle; IVS – interventricular septum; LA – left atrium; LV – left ventricle’ PA – pulmonary artery; RA – right atrium; RV – right ventricle; asterisks – denote lumenized vasculature. **G-H,** β-gal activity in P1 postnatal lungs from (A) *Dll4*^*lacZ/+*^ or (B) *Dll4-BAC-nlacZ* mice. **G1-G2** and **H1-H2** show representative wholemount lungs from *Dll4*^*lacZ/+*^ and *Dll4-BAC-nlacZ* mice, respectively, from ventral and dorsal views. **G3** and **H3** show β-gal activity in a representative cross-section through the lungs, which is magnified accordingly in panels **G4** and **H4. I-J,** β-gal activity in P5 postnatal lungs from (C) *Dll4*^*lacZ/+*^ or (D) *Dll4-BAC-nlacZ* mice. **I1-I2** and **J1-D2** show representative wholemount lungs from *Dll4*^*lacZ/+*^ and *Dll4-BAC-nlacZ* mice, respectively, from ventral and dorsal views. **I3** and **J3** show β-gal activity in a representative cross-section through the lungs, which is magnified accordingly in panels **I4** and **J4. K-L,** β-gal activity in adult lungs from (E) *Dll4*^*lacZ/+*^ or (F) *Dll4-BAC-nlacZ* mice. **K1-K2** and **L1-L2** show representative wholemount lungs from *Dll4*^*lacZ/+*^ and *Dll4-BAC-nlacZ* mice, respectively, from ventral and dorsal views. **K3** and **L3** show β-gal activity in a representative cross-section through the lungs, which is magnified accordingly in panels **K4** and **L4.** In both lines, and at all stages, β-gal activity appears to be confined to the endothelium. Abbreviations: D – dorsal; e – endothelium; sm – smooth muscle; L – left; R – right; V - ventral

β-gal was present within the trachea in the postnatal and adult lung, at all stages examined, in both lines. At P1 and P5, the endothelium of the small, medium, and large caliber vessels, but not the smooth muscle or airway epithelium, displayed *lacZ* expression in both *Dll4*^*lacZ/+*^ and BAC animals. The alveoli were also β-gal positive, with expression in the capillary endothelium (Fig. 7G1-H4). This expression pattern perdured in adults, with the only notable difference between the two lines being the extent of activity within the alveoli (Fig. 7K1-L4).

In the postnatal retina, some notable differences in expression were observed between the two lines. At P1 (Fig. 8A1-B3), *Dll4*^*lacZ/+*^ expression within the vasculature was absent (Fig. 8A1-A3), but signal was present (though minimal) in the vessels of *Dll4-BAC-nlacZ* animals (Fig. 8B1-B3). This difference was more pronounced at P5, where signal was virtually absent within the vasculature of *Dll4*^*lacZ/+*^ animals (Fig. 8C1-C3), but strong in *Dll4-BAC-nlacZ* animals (Fig. 8D1-D3). By P7, *lacZ* expression was more comparable between the reporter lines, though still diminished in *Dll4*^*lacZ/+*^ mice compared to the BAC reporter line (Fig. 8E1-F3). By adulthood, no gross differences were observed in staining between the two alleles (Fig. 8G1-H3), with labelling throughout the retinal vasculature. In both lines X-gal signal was detectable in the tissue underlying the surface vasculature (presumably astrocytes) at all stages examined, although this was greatly diminished in the adult retina. Vascular signal in either genotype appeared arterial-specific, and was present within the capillary vasculature in the adult retina (Fig. 8G1-H3). To determine if *lacZ* expression was restricted to arteries, adult retinas were immunostained for β-gal, the pan-endothelial marker isolectin B4, and the smooth muscle cell marker SMA (as smooth muscle cells are associated with arteries). In *Dll4-BAC-nlacZ* retinas, colocalization was observed between all three markers (β-gal, isolectin, and SMA) (Fig. 8I1-I6), suggesting that reporter expression was indeed restricted to the arterial and capillary endothelium. β-gal immunohistochemistry on adult retinas from knockin reporter animals failed to yield interpretable results (Fig. S1), regardless of fixation method, primary antibody concentration, or length of antibody incubation. This is perhaps attributable to diminished *Dll4* expression, which is consistent with tissues processed for X-gal staining, as knockin tissue required longer incubation times to achieve adequate signal compared to BAC reporter samples. Differences in clarity were prominent at all stages examined, with *Dll4-BAC-nlacZ* retinas displaying better cellular resolution.

**Figure 8:**
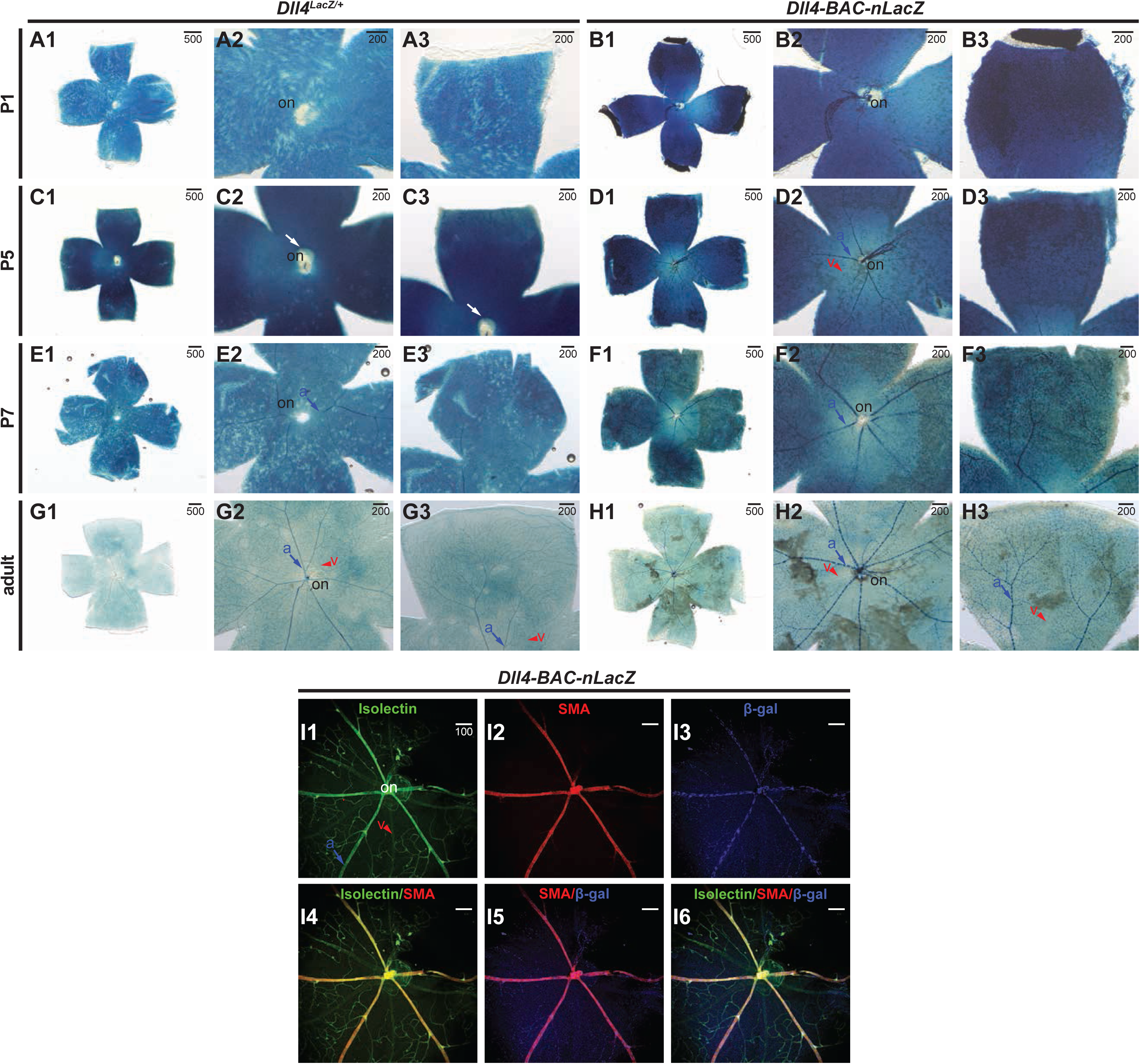
Comparative *Dll4* expression in postnatal and adult retinas. **A-B,** β-gal activity in P1 postnatal retinas from (A) *Dll4*^*lacZ/+*^ or (B) *Dll4-BAC-nlacZ* mice. Both lines are clearly active in non-endothelial cell types within the retina at this stage. **C-D,** β-gal activity in P5 postnatal retinas from (C) *Dll4*^*lacZ/+*^ or (D) *Dll4-BAC-nlacZ* mice. Dll4-BAC-nlacZ is more active than the endogenous reporter in the developing arterial endothelium. **E-F,** β-gal activity is detected in the arterial and capillary endothelium in both (E) *Dll4*^*lacZ/+*^ and (F) *Dll4-BAC-nlacZ* retinas at P7. **G-H,** *Dll4*^*lacZ/+*^ is enriched in arteries and capillary vessels (and weakly detected in veins), while *Dll4-BAC-nlacZ* has no activity within the veins in the adult retina. **I-J,** Immunohistochemistry and indirect immunofluorescent detection of isolectin (I1), smooth muscle actin (I2), β-gal (I3), and merged (I4-I6) images from representative *Dll4-BAC-nlacZ* adult retinas. Abbreviations: a – artery; on – optic nerve; v – vein.

## Discussion

In the present study, we generated a novel *Dll4* reporter and compared its expression to a commonly used *Dll4* knockout / *lacZ* knockin line (Duarte et al., 2004). This novel tool avoids the confounding variable of haploinsufficiency associated with previous *Dll4* knockin reporter alleles. In addition, this new line allows for increased resolution of *Dll4* expression due to the presence of a nuclear-localized reporter, and it generally recapitulates *Dll4* expression in the embryo and adult. Our novel BAC reporter, however, does exhibit important differences in expression compared to the knockin model used for our comparative analyses, and these points will be discussed below on a tissue by tissue basis.

The results of our studies with the BAC reporter during embryogenesis mirror previously published data examining *Dll4* expression by *in situ* hybridization (Chong et al., 2011; Mailhos et al., 2001; Shutter et al., 2000; Villa et al., 2001), and are generally concordant with studies of *Dll4* knockin reporter alleles (Duarte et al., 2004; Gale et al., 2004; Wythe et al., 2013). Specifically, *lacZ* was detected within aortic progenitor cells at E7.75, and the endothelial-specific, arterial-restricted activity of β-gal was maintained in the main axial vessels through the formation of a functional dorsal aorta at E10.5 (Chong et al., 2011; Wythe et al., 2013). Expression in other arterial beds, such as the internal carotid artery, intersegmental arteries, and umbilical artery were all equivalent to the knockin reporter through age E10.5. Thereafter, both the knockin and BAC lines retained their endothelial, arterial-specificity within the skin of the embryonic E14.5 forelimb. Collectively, these results confirm that in the developing embryo the BAC reporter functions as a accurate indicator for arterial endothelial identity, as well as *Dll4* expression, free from the arteriovenous patterning defects endemic to *Dll4* knockin reporter alleles.

Notch/Rbpj-k signaling represses mesodermal adoption of a myocardial cell fate within in *Xenopous* (Rones et al., 2000), consistent with *in vitro* (Schroeder, 2003 #71;) and in vivo murine studies (Watanabe et al., 2006), as well as work in *Drosophila* (Han and Bodmer, 2003; Rones et al., 2000). Of the Notch signaling machinery, *Dll1*, *Notch1* and *Notch4* are also transcribed in the early endocardium (Del Amo et al., 1992; Grego-Bessa et al., 2007; Uyttendaele et al., 1996), whereas *Notch2* and *Jag2* are expressed in the myocardium at later stages (Loomes et al., 1999; McCright et al., 2002). Within the heart, β-gal was first detected in both lines in the presumptive endocardium of the early cardiac crescent (∼E7.5-E7.75), mimicking endogenous *Dll4* transcripts (Mailhos et al., 2001; Shutter et al., 2000; Wythe et al., 2013). Expression throughout the endocardium was maintained through embryonic turning to age E9.5. Prior work suggested that *Dll4* is preferentially transcribed within the endocardium at the base of the cardiac trabeculae at E9.5, while *Notch1* is expressed uniformly throughout the endocardium (Grego-Bessa et al., 2007). Herein, at E9.5 both reporters showed a non-uniform, salt and pepper expression pattern within the ventricular endocardium (potentially favoring the base of the trabeculae), suggestive of lateral inhibition (or induction), a classical mechanism of juxtacrine Notch signaling (Lewis, 1998). This expression agrees with reports of Notch activation in the endocardium at E9.5 (D’Amato et al., 2016; Grego-Bessa et al., 2007). β-gal signal was robust in the outflow tract and atrioventricular canal at E9.5 in both lines. At E10.5, in the knockin and BAC reporter β-gal appeared more evenly distributed throughout the ventricular and atrial endocardium, and was evident in the distal—but not proximal—outflow tract. Critically, pan-endothelial deletion of *Notch1* or *Rbpj-k* (via *Tie2:Cre*) yields severe trabeculation defects (Grego-Bessa et al., 2007), presumably by disrupting critical endocardial to myocardial signaling networks (Meyer and Birchmeier, 1995). Recent work from the de La Pompa lab suggested that Manic Fringe (Mfng) activity within the endocardium favors Dll4-mediated activation of Notch1 over myocardial-derived Jag1 in the early heart, and subsequent downregulation of *Dll4* and *Mfng* coupled with upregulation of *Jag1*/*Jag2* favors myocardial cell activation of endocardial Notch1 later in heart development (D’Amato et al., 2016). Myocardial deletion of *Mindbomb* (*Mib*), which encodes an E3 ubiquitin ligase required for Notch ligand function, generates left ventricular non-compaction (due to lack of trabeculation) (Luxan et al., 2013). Intriguingly, *Jag1/2* compound myocardial-specific knockouts fail to recapitulate all phenotypic defects seen in *cTnT-Cre*;*Mib*^*flox/flox*^ animals, suggesting another Notch ligand may be active within the myocardium. Reporter activity was detected in the myocardium at E18.5 (as early as E14.5) in both the *Dll4* knockin and BAC lines. The biological significance of *Dll4* expression within the developing myocardium remains to be determined. β-gal was also observed in the adult myocardium in both reporters, suggesting a possible role in adult myocyte function. It would be interesting to determine if Dll4-Notch1 interactions affect myocardial behavior in a pathologic setting.

In the developing and adult lung, β-gal was evident within the trachea and endothelium of large and small caliber vessels, as well as the capillaries surrounding the alveoli. Notch signaling in the trachea, as with many other tissues, is known to regulate cell fate decisions, with Notch gain-and loss-of-function manipulations resulting in the failure of proper tracheal branching in *Drosophila* (Llimargas, 1999; Steneberg et al., 1999). Furthermore, alveologenesis requires Notch signaling in the mouse lung epithelium (Tsao et al., 2016). *Dll4* also has a physiological role in allergic inflammatory responses in the airway (Huang et al., 2017). It is possible that Dll4 presented on endothelial cells activates Notch receptors on adjacent cell types to regulate patterning of the trachea and lungs, a notion supported by reporter expression in adult pulmonary tissues in both the BAC and knockin lines.

At E9.5 and E10.5, the arterial cranial vasculature was labelled in both lines (Fig. 1). At E12.5, β-gal was evident in the arterial vasculature of brains from both lines, although MCA labeling was diminished in knockin animals. Previous work reported that loss of *Dll4* delayed MCA formation and resulted in hyperbranching (Cristofaro et al., 2013), defects that were observed here as well (Fig. 4). However, knockin brains were often smaller at this stage, and general developmental delay may have caused this defect. The BAC line did not present this phenotype. Brains between the two genotypes were comparable in size at E14.5 and E18.5, but the caliber of vessels feeding into the circle of Willis, as well as the basilar artery, appeared larger in the BAC animals than in the knockins at E18.5. In contrast to previous reports suggesting that *Dll4* is excluded from large caliber arteries at later stages (Benedito and Duarte, 2005; Gale et al., 2004), expression was evident in the major cranial arteries of the postnatal and adult brain in both reporters (Fig. 6). Collaterals were well-labelled in both lines, but potentially more obvious in the BAC line. Dll4-Notch signaling is essential for embryonic vascular development (Duarte et al., 2004; Gale et al., 2004; Krebs et al., 2004; Krebs et al., 2000; Swiatek et al., 1994), and the continued expression of *Dll4* in the brain may suggest a role in regulating angiogenesis. Indeed, deletion of *Rbpj-k* at birth leads to arteriovenous malformations and increased vascular density in the brain at P14, followed shortly thereafter by lethality (Nielsen et al., 2014). By comparison, *Rbpj-k* loss in the adult mouse produced a mild phenotype (Nielsen et al., 2014). However, Dll4-Notch signaling has been suggested to modulate angiogenic responses in the brain after ischemic injury (Cristofaro et al., 2013). Given Notch’s role in regulating neurogenesis from development through adulthood (de la Pompa et al., 1997; Imayoshi and Kageyama, 2011; Imayoshi et al., 2010), it would be interesting to discern what role(s), if any, endothelial Dll4-Notch signaling has in embryonic and adult neurogenesis.

In the periphery, nerves and arteries regularly align with one another (Mukouyama et al., 2002), as nerves induce arteriogenesis by secreting VEGF and CXLC12 (Li et al., 2013; Mukouyama et al., 2005). β-gal^+^ cells in the forelimb skin of BAC embryos align with Tuj1^+^ nerves, are CD31^+^, and are encapsulated by SMA^+^ cells, demonstrating they are bona fide arterial endothelial cells (Fig. 3). Notably, endogenous Dll4 displayed a pattern identical to that of the reporters (Fig. 3). As such, it would be interesting to determine if endothelial Dll4 plays a role in nerve-vessel alignment.

Reporter activity within the postnatal and adult retina agree with published reports showing *Dll4* transcript and protein expression in the endothelium at P3 (Crist et al., 2017; Hofmann and Luisa Iruela-Arispe, 2007), supporting the concept that *Dll4* heterozygosity in the knockin delays reporter expression in the postnatal retina. Vascular *lacZ* expression was virtually absent until P7 in the knockin, but BAC reporter activity was detectable at low levels in the center of the retina, near the optic nerve, as early as P1. The ultimate impact of delayed *Dll4* expression in the postnatal mutant eye (Suchting et al., 2007) may be inconsequential, as vascular patterning in the adult retina was grossly indistinguishable between the two reporter lines (Fig. 8 G1-H3). Surprisingly, arterial-specific deletion of *Dll4* or *Rbpj-k* at P10, after the major vascular network has been patterned, does not generate profound vascular remodeling defects in the retina by P28 (nor does deletion at P2 affect vascular structure at P15) (Ehling et al., 2013). However, pan-endothelial deletion of either gene induced vascular defects (Ehling et al., 2013), suggesting a role for Notch signaling in the capillary endothelium and venous tissue, potentially in agreement with reports of capillary and venous *Dll4* expression in the retina (Crist et al., 2017; Ehling et al., 2013).

Notably, the BAC allele tended to reveal *Dll4* expression at earlier time points compared to the knockin. This may result from increased *lacZ* expression in the BAC reporter due to transgene copy number, or it may be attributable to normal levels of Notch signaling (unlike in the knockin mutants). In the postnatal retina delayed *lacZ* expression in the knockin, but not the BAC, suggests that *Dll4* heterozygosity, even in a genetic background meant to mitigate the effects of its haploinsufficiency (e.g. CD1 or FVB), still negatively influences reporter expression. Additionally, the knockin line required significantly longer (by several hours) incubation times for adequate visualization of reporter activity in all tissues examined. In the BAC line, 30 minutes was usually more than adequate for robust visualization of β-gal staining and resolution of single endothelial cells. Overall, the BAC reporter generates a more representative pattern of *Dll4* expression than the heterozygous knockin reporter.

Multiple studies have suggested that *Foxc1/2*, as well as *β-catenin*, mediate the transcriptional induction of *Dll4* within the embryonic endothelium (Corada et al., 2010; Hayashi and Kume, 2008; Seo et al., 2006). However, previous work demonstrated that endothelial-specific deletion of *β-catenin* failed to affect establishment of arteriovenous identity or alter *Dll4* expression in the early mouse embryo (Wythe et al., 2013). Additionally, the genomic region (5’ to the transcriptional start site (TSS) of murine *Dll4*) shown to bind these same transcription factors failed to drive reporter activity *in vivo* (Wythe et al., 2013), suggesting alternative transcriptional regulators of *Dll4*. Indeed, we, and others, identified enhancers within the third intron of *Dll4*, as well as several kilobases upstream of the TSS (-10, and -12, respectively) (Luo et al., 2012; Sacilotto et al., 2013; Wythe et al., 2013). The targeting vector that generated the *Dll4*^*lacZ/+*^ allele utilized in this study retained the intron 3 enhancer (Duarte et al., 2004). However, another *Dll4* knockin/knockout reporter mouse with arterial *lacZ* expression (Gale et al., 2004) replaced the entire *Dll4* locus, suggesting that this region is dispensable for endothelial expression of *Dll4* (we are not aware of any study directly comparing β-gal activity between these two mutant lines). Nonetheless, the 81-kb region spanned by the BAC reporter contains each of these *in vivo* validated genomic elements sufficient to drive *Dll4* arterial expression. Future deletion of these, and other conserved regulatory elements, will determine the necessity of these putative enhancers.

In the absence of genetic reporter models, studies must rely on molecular and biochemical methods to assess gene expression *in vivo*. *In situ* hybridization using nucleic acid probes is a common method for analyzing gene expression patterns. This technique, however, relies on the quality and fidelity of the probes used to detect mRNA transcripts of interest and, as such, can exhibit high variability from one probe to the next. Furthermore, the utility of this method is limited by riboprobe penetration and cellular resolution in wholemount tissue. Immunostaining presents similar obstacles regarding variability, as often multiple commercial antibodies exist for the same antigen, and several are derived from finite sources (e.g. polyclonal). Our mouse model provides a simple, robust, renewable, and reliable alternative approach to visualize *Dll4* expression *in vivo*. Furthermore, this *Dll4* mouse line can be combined with other alleles and genetic backgrounds to elucidate epistatic interactions, without the requirement of having to assess its expression on a confounding heterozygous background, as is the case with current *Dll4* reporters. Furthermore, as demonstrated herein, this allele is well-suited for immunohistochemistry studies due to the restricted nuclear localization of the antigen. Going forward, this new mouse line can be used to determine how *Dll4* expression changes in response to gain - or loss-of-function gene manipulations, and whether levels of *Dll4* are changed in response to injury, disease, or drug treatment. Collectively, this novel *Dll4-BAC-nlacZ* reporter mouse line will prove a valuable tool in deciphering the mechanisms underlying Notch signaling, and will provide researchers with a useful reagent for investigating Dll4-associated mechanisms.

## Materials and Methods

### Mouse experiments

All mouse protocols were approved by the Institutional Animal Care and Use Committee (IACUC) at Baylor College of Medicine and at UCSF. For all experiments, noon on the day a plug was discovered was considered embryonic day 0.5.

### Cloning and Recombineering

To create a transgenic reporter insertion at the ATG of *Dll4*, we purchased a mouse BAC clone, bMQ132J23, from the AB2.2 ES cell DNA (129S7/SvEv Brd-Hprt b-m2) derived bMQ library (Cox et al, 2005) (Source Biosciences). This clone spans nucleotides 119,293,232-119,374,281 on chromosome 2. DNA was purified from the BAC clone and transformed by electroporation into SW102 bacteria (for subsequent GalK manipulation of DNA). Bacteria were plated on LB chloramphenicol plates and the resulting colonies were screened by PCR for the 5’ and 3’ ends of the BAC to confirm successful transmission. After confirmation, the *loxP511* and *loxP* sites were removed from the BAC. Briefly, *loxP511* and *loxP* GalK replacement fragments were created by amplifying the EM7-GalK open reading frame through PCR using two oligos with 50 bp homology arms to pBACe2.6.

**loxP511 GalK ins FWD**: CGT AAG CGG GGC ACA TTT CAT TAC CTC TTT CTC CGC ACC CGA CAT AGA TAC CTG TTG ACA ATT AAT CAT CGG CA

**loxP511 GalK ins REV**: CGG GGC ATG ACT ATT GGC GCG CCG GAT CGA TCC TTA ATT AAG TCT ACT AGT CAG CAC TGT CCT GCT CCT T

**loxP GalK ins FWD**: CTT ATC GAT AAG CTG TCA AAC ATG AGA ATT GAT CCG GAA CCC TTA ATC CTG TTG ACA ATT AAT CAT CGG CA

**loxP GalK ins REV**: CCG ATG CAA GTG TGT CGC TGT CGA CGG TGA CCC TAT AGT CGA GGG ACC TAT CAG CAC TGT CCT GCT CCT T

Each GalK cassette was sequentially inserted and then replaced by a single, 100 bp oligo that lacks the original *loxP511* or *loxP* sequences, but contains the original plasmid backbone sequence, effectively deleting the *loxP511* and *loxP* sequences and leaving no scar.

***loxP511* replacement oligos**:

**FWD**: CGT AAG CGG GGC ACA TTT CAT TAC CTC TTT CTC CGC ACC CGA CAT AGA TAC TAG TAG ACT TAA TTA AGG ATC GAT CCG GCG CGC CAA TAG TCA TGC CCC G

**REV**: CGG GGC ATG ACT ATT GGC GCG CCG GAT CGA TCC TTA ATT AAG TCT ACT AGT ATC TAT GTC GGG TGC GGA GAA AGA GGT AAT GAA ATG TGC CCC GCT TAC G

***loxP* replacement oligos**:

**FWD**: CTT ATC GAT GAT AAG CTG TCA AAC ATG AGA ATT GAT CCG GAA CCC TTA ATT AGG TCC CTC GAC TAT AGG GTC ACC GTC GAC AGC GAC ACA CTT GCA TCG G

**REV**: CCG ATG CAA GTG TGT CGC TGT CGA CGG TGA CCC TAT AGT CGA GGG ACC TAA TTA AGG GTT CCG GAT CAA TTC TCA TGT TTG ACA GCT TAT CAT CGA TAA G

PCR genotyping for successful replacement of *loxP* sites was performed with the following primer pairs:

***loxP511*-FWD**: GGC AGT TAT TGG TGC CCT TA

***loxP511*-REV**: TTC AAC CCA GTC AGC TCC TT

expected size=353 bp

***loxP*-FWD**: TAG TGA CTG GCG ATC CTG TC

***loxP*-REV**: AAC ATT TTG CGC ACG GTT AT

expected size=396 bp

At this point, the resulting plasmid was referred to as Δ*loxP*-*Dll4*-BAC. Next, a 5’ homology arm to murine *Dll4* was amplified by PCR with the following primers (pGalK homology in lowercase, unique restriction sites are in italics, *Dll4* homology underlined in capitals):

**FWD**: accgggccccccctcgag*GTCGAC*<u>ACTGTAGCCACTAGAGGCCTG</u>

**REV(EcoRV)**: tgtcaacaggaattc*GATATC*<u>CATCCCTTGGGGTGTCCTCTCCAC</u>

The resulting fragment was cloned via cold fusion (SBI) into the digested and purified pGalk vector 5’ to the EM7-GalK cassette. After identification of a positive clone and confirmation by DNA sequencing, a 3’ *Dll4* homology arm was amplified and then inserted 3’ to the GalK cassette into the SpeI and NotI sites using the following primers:

**FWD**: gacagtgctgaggatcc*ACTAGT*<u>ACGCCTGCGTCCCGGAGCGCC</u>

**REV (NotI)**: tccaccgcggtg*GCGGCCGC*<u>ACCGGCGTGGAGACATTGCCAAAGG</u>

The *Dll4* 5’ 3’ arm GalK vector was digested, the homology arm Galk fragment purified, and then electroporated into SW102 Δ*loxP*-*Dll4*-BAC bacteria for positive selection on M63 + galactose plates to isolate a Δ*loxP*-*Dll4*-BAC-GalK clone. Concurrently, a codon-optimized nls-lacZ (from Invivogen’s pWhere plasmid) was subcloned between the same 5’ and 3’ *Dll4* homology arms, into the EcoRI site (5’) and BamHI site (3’). After confirmation by sequencing, *Dll4* 5’ 3’ arm nls-lacZ-pA, was digested with SalI and NotI to release the targeting fragment, and after purification this element was transformed into electro-competent Δ*loxP*-*Dll4*-BAC SW102-GalK bacteria and subjected to negative selection on M63 plates + DOG. The colonies were screened by PCR and the resulting construct, Δ*loxP*-*Dll4*-BAC-nlacZ, was confirmed by DNA sequencing.

### Generation of transgenic mice

The Δ*loxP*-*Dll4*-nlacZ-BAC DNA was purified using the BAC 100 prep kit (Nucleobond) and digested with PI-SceI to linearize the BAC for more efficient transgenesis. A portion was inspected by pulse field electrophoresis to confirm the correct restriction pattern, then the remainder was dialyzed (Spectra/Por Micro DispoDialyzer; 8,000 Da MWCO, 100 μL) to embryo water (Sigma, # W1503), and used for pronuclear injection. Injection of transgenic fragments was performed at the Gladstone Institute. *Dll4*-BAC-nlacZ^#4316^ is a weaker, but consistent founder line and *Dll4*-BAC-nlacZ^#4336^ is a strong expresser.

### Genotyping and mice used

*Dll4*^*lacZ/+*^ (*Dll4*^*tmJrt*^) (Duarte et al., 2004) cryopreserved embryos were purchased from the Canadian Mouse Mutant Repository (CMMR) and implanted into CD31 females. Genotyping for all alleles was performed by PCR.

**JDW 10 (Dll4 FWD)**: 5’-GGGGAATCAGCTTTTCAGGAA

**JDW 11 (Dll4 REV)**: 5’-CGAACTCCTGCAGCCGCAGCT

**JDW 12 (KO REV)**: 5’-ACGACGTTGTAATACGAC

The knockout allele generates a 110 bp band, while the WT allele results in a 300 bp band (these primers were not multiplexed).

**JDW 138 (*Dll4* 5’ UTR)**: 5’-CTC TGG AGC AAG CAG GTT TC

**JDW 139 (nlacZ ORF)**: 5’-TTG AAG GTC AGG CTG TAG CA

The presence of the transgene generates a 750 bp product.

### lacZ Staining

Embryos were harvested at timed intervals. A vaginal plug in the morning indicated embryonic day 0.5. Appropriately timed-mated, pregnant dams were euthanized by CO_2_, and embryos were carefully dissected away from all internal membranes into cold 1X PBS. For whole-mount processing, embryos were processed as previously described (Wythe et al., 2013). Briefly, following dissection, embryos or their dissected organs (brain, heart, or lungs) were fixed with a formaldehyde/glutaraldehyde solution in 1X PBS (2% formaldehyde, 0.2% glutaraldehyde, 0.02% sodium deoxycholate, 0.01% NP-40). For embryos and organs ≤E8.5, fixation time was 5 minutes. For embryos and organs ≤E10.5, fixation time was 10 minutes. For all older embryos and organs, fixation time was 15-20 minutes.

Following fixation, tissues were rinsed briefly in 1X PBS and embryos were placed in permeabilization solution (1X PBS, 0.02% sodium deoxycholate, 0.01% NP-40). Embryos and tissue ≥E10.5 were incubated in permealibilization buffer overnight to allow for sufficient penetration of staining. After permeabilization, embryos were incubated at 37°C in freshly-made, 0.22 µm-filtered X-gal staining solution made in permeabilization buffer (5mM potassium ferricyanide, 5mM potassium ferrocyanide, 2mM MgCl_2_, 1 mg/mL X-gal). Whole embryos and organs were incubated in this solution between 3-4 hours for *Dll4-BAC-nlacZ* animals, though often 6-8 hours for *Dll4*^*lacZ/+*^ animals to acquire similar levels in staining. Embryos and organs were then rinsed briefly twice in permeabilization buffer to remove residual staining solution, followed by a longer 20-minute wash. This was followed by post-fixation overnight in 4% PFA at 4°C. The following day, PFA was removed and embryos and organs were washed twice for 10 minutes each wash in PBST (1X PBS, 0.1% Tween-20). Tissues were then subjected to a serial dehydration with methanol (25% MeOH/PBST, 50% MeOH/PBST, 75% MeOH/PBST, and finally 3 washes of 100% MeOH for 10 minutes each wash). Lastly, embryos and organs were washed with 5% H_2_O_2_/ 95% MeOH for 1 hour at room temperature. Larger embryos (≥E12.5) were further washed with 7.5% H_2_O_2_/MeOH for 15 minutes if any yellowing of the tissue was still present. Tissues were then serially rehydrated in PBST, then stored in 4% PFA until imaging or further processing.

For embryonic and early postnatal brain sections, brains were dissected and rinsed in 1X PBS, followed by overnight fixation in 2% PFA at 4°C. For adult brains, mice were transcardially perfused with 1X PBS followed by 4% PFA before brains were removed. Brains were then transferred into serial sucrose/PBS solutions (10%, 20%, and 30%) and then frozen in optimal cutting temperature (OCT) compound and stored at -80°C. Adult and early postnatal brains were cryosectioned at 40 µm and placed in 2% PFA for 15 minutes (free-floating). Sections were then rinsed briefly in 1X PBS before being washed twice in permeabilization buffer for 10 minutes each wash. The tissue was then stained at 37°C in X-gal staining solution for 3 hours for *Dll4-BAC-nlacZ* animals, and 4-5 hours for *Dll4*^*lacZ/+*^ animals. After staining, sections were briefly rinsed several times to eliminate staining solution and mounted using Fluoromount-G mounting media (SouthernBiotech, 0100-01). For embryonic brains, tissue was processed similarly, with the exception that sections were placed directly onto slides after sectioning, rather than using a free-floating method. For embryonic and early postnatal heart and lung sections, tissues were harvested and fixed for 2 hours in 2% PFA at 4°C. Tissues were subsequently transferred into serial sucrose/PBS solutions (10%, 20%, and 30%) and then frozen in optimal cutting temperature (OCT) compound and stored at -80°C. Cryosections were taken at 10 µm and mounted directly to glass slides for processing. X-gal staining was performed as previously stated, but were further processed afterwards for eosin (*Dll4-BAC-nlacZ*) or nuclear fast red (*Dll4*^*lacZ/+*^) staining. Slides were submerged in eosin solution (Thermo Scientific, #7111) for 3 minutes before being washed 2X for 3 minutes each wash in tap water. Slides were then dipped 3X for 30 seconds in 100% EtOH, followed by 3X for 1 minute in xylene. For nuclear fast red staining (Vector Labs, H-3403), slides were submerged for 15 minutes, followed by identical washes in tap water, EtOH, and xylene. Slides were then mounted using Entellan New (Millipore, #107961). For adult heart and lungs, mice were first transcardially perfused with 1X PBS and 2% PFA before identical post-fixation as stated above. Adult lungs were also first infused with 1% low melting point agarose and allowed to solidify prior to post-fixation and processing.

### Immunohistochemistry

IHC performed on embryonic limb skin was performed according to Mukouyama et al (Mukouyama et al., 2012). Briefly, forelimbs from E14.5 embryos were removed in ice-cold 1X PBS and subsequently transferred to 4% PFA at 4°C overnight. On the next day, tissue was dehydrated in 100% MeOH and stored at -20°C. Forearm limb skin was gently removed from the underlying tissues and placed in 100% MeOH. Samples were then rehydrated by transferring them into 75%/50%/25% MeOH/1X PBST (1X PBS with 0.2% Trition X-100) for 5 minutes each step. Samples were then washed twice for 5 minutes in 1X PBST before putting the tissue in filter-sterilized blocking solution (10% horse serum, 0.5% Triton-X, 1X PBS) for 2 hours at room temperature. Blocking solution was then removed and primary antibodies prepared in the same blocking solution were added and left overnight to shake gently at 4°C. Skin samples were then washed in blocking buffer 5 times for 10 minutes each wash before adding secondary antibodies diluted in blocking buffer, and allowed to incubate for 1 hour at room temperature. Tissues were then washed again in blocking buffer 5 times for 10 minutes each wash and mounted on glass slides using Fluoromount-G mounting media (SouthernBiotech, 0100-01) and imaged.

For retinas, adult eyes were removed and placed in 4% PFA overnight at 4°C. Retinas were then removed the following day and placed in retina blocking buffer (1% BSA, 0.3% Triton X-100, 1X PBS) overnight at 4°C. Retinas were then placed into Pblec solution (1 mM MgCl_2_, 1 mM CaCl_2_, 0.1mM MnCl_2_, 1% Triton X-100, 1X PBS) and washed 3 times for 20 minutes each wash at room temperature. Primary antibodies were prepared in Pblec solution, added to the retinas, and allowed to incubate overnight at 4°C with gentle shaking. The next day, retinas were washed 5X for 5 minutes each wash with retina blocking buffer diluted 1:1 in 1X PBS. Secondary antibodies were then added for 2 hours at rooms temperature followed by 5, 10 minute washes with retina blocking buffer diluted 1:1 in 1X PBS. Retinas were then mounted on glass slides using Fluoromount-G mounting media and imaged. Primary antibodies used were: goat anti-Dll4 (1:100) (R&D Systems, AF1389), rat anti-CD31 (1:200) (BD Pharmingen, 550274), hamster anti-Podoplanin (1:100) (Hybridoma Bank, 8.1.1), mouse anti-Tuj1 (1:500) (Covance, MMS-435P), rabbit anti-β-galactosidase (1:1000) (MP Biomedical, 55976), biotinylated Griffonia Simplicifolia Lectin I isolectin B4 (1:50) (Vector Labs, B-1205), and mouse anti-actin, alpha-smooth muscle FITC (1:100) (Sigma, F3777). All secondary antibodies (1:200) (Life Technologies) were used at at room temperature and included the following antibodies depending on primary combinations used: donkey anti-goat 488, goat anti-rat 594, and goat anti-rabbit 647. Anti-actin, alpha-smooth muscle was conjugated with a FITC fluorescein molecule and was always added during the secondary antibody step for 1-2 hours. For visualization of the isolectin primary antibody, DyLight 594 Streptavidin (1:100) (Vector Labs, SA-5594) secondary antibody was added for 1-2 hours. For IHC that required goat anti-Dll4 primary antibody, secondary antibodies were added sequentially to prevent potential cross-reactivity. Donkey anti-goat secondary was added first for one hour at room temperature, followed by 4 brief washes with copious amounts of blocking solution, followed by 3, 10 minute washes with blocking solution. Afterwards, goat anti-rabbit and goat anti-rat secondary antibodies were added for one hour, washed, and mounted.

### Imaging

Skin and retinas processed for immunohistochemistry were imaged using a Leica TCS SPE confocal microscope with a 10X objective. All other tissues and embryos were imaged using a Zeiss Axio Zoom.V16 microscope and processed using ZenPro software and Adobe Photoshop. All images were assembled using Adobe Illustrator.

## Acknowledgements

The authors thank the Arenkiel laboratory at the NRI at BCM for use of their confocal microscope, and Karen Berman de Ruiz for superb mouse colony care and maintenance.

### Competing Interests

No competing interests to declare.

### Author Contributions

JDW was responsible for the conception, design, execution, supervision and interpretation of experiments. JDW and AMH wrote the manuscript. AMH was involved in the execution and analysis of experiments. AMR performed experiments. JDW and WPD generated the BAC reporter construct. WPD edited the manuscript. SPM was involved in the design and analysis of experiments, and edited the manuscript. BB provided reagents and financial support and edited the manuscript. All authors revised the manuscript and consented to its contents.

### Funding

AMH was supported by the NIH (2T32HL007676). WPD was a California Institute for Regenerative Medicine Clinical Scholar (TG2-01160) and supported by the NIH (5T32-HL007731-20). SPM is supported by the National Institutes of Health National Instittue of Neurological Disorders and Stroke (R01 NS094280 and R01 NS096186) BG. was funded by the National Institutes of Health National Heart, Lung, and Blood Institute (NHLBI) (Bench to Bassinet Program UM1 HL098179; P01 HL089707) and by William H. Younger, Jr. JDW is supported by institutional startup funds from the CVRI at Baylor College of Medicine, the Caroline Wiess Law Fund for Research in Molecular Medicine, the Curtis Hankamer Basic Research Fund, and the ARCO Foundation Young Teacher-Investigator Award. Work within the Wythe lab is supported by the AHA (12SDG12060353 and 16GRNT31330023).

## Supplementary Material

**Figure S1:**
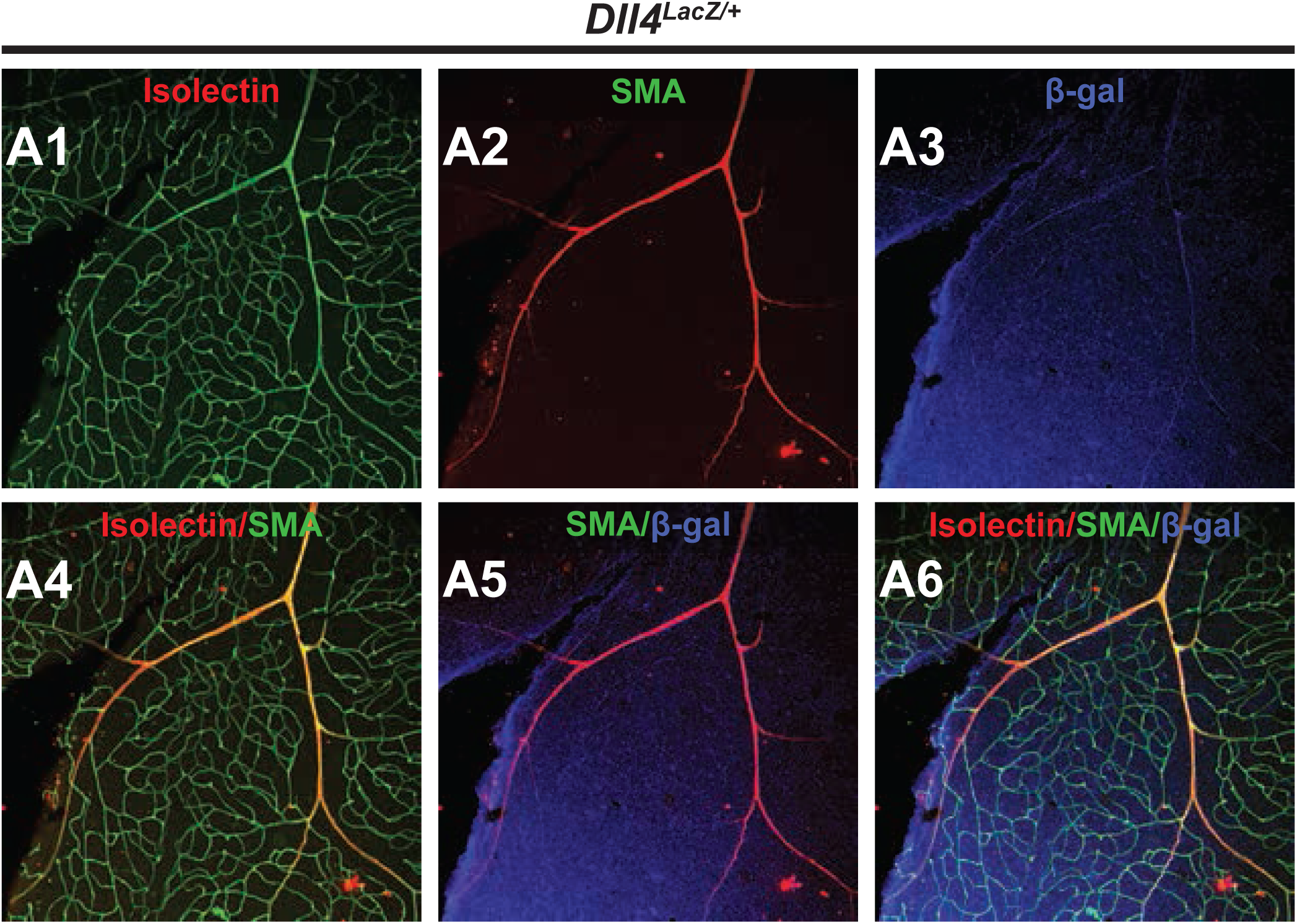
Comparative *Dll4* expression in postnatal and adult retinas. **A1-A6,** Immunohistochemistry and indirect immunofluorescent detection of isolectin B4 (A1), smooth muscle actin (SMA) (A2), β-gal (A3), and merged (A4-A6) images from representative *Dll4*^*lacZ/+*^ adult retinas.

